# snpXplorer: a web application to explore human SNP-associations and annotate SNP-sets

**DOI:** 10.1101/2020.11.11.377879

**Authors:** Niccolo Tesi, Sven van der Lee, Marc Hulsman, Henne Holstege, Marcel J.T. Reinders

## Abstract

Genetic association studies are frequently used to study the genetic basis of numerous human phenotypes. However, the rapid interrogation of how well a certain genomic region associates across traits as well as the interpretation of genetic associations is often complex and requires the integration of multiple sources of annotation, which involves advanced bioinformatic skills. We developed *snpXplorer*, an easy-to-use web-server application for exploring Single Nucleotide Polymorphisms (SNP) association statistics and to functionally annotate sets of SNPs. *snpXplorer* can superimpose association statistics from multiple studies, and displays regional information including SNP associations, structural variations, recombination rates, eQTL, linkage disequilibrium patterns, genes and gene-expressions per tissue. By overlaying multiple GWAS studies, *snpXplorer* can be used to compare levels of association across different traits, which may help the interpretation of variant consequences. Given a list of SNPs, *snpXplorer* can also be used to perform variant-to-gene mapping and gene-set enrichment analysis to identify molecular pathways that are overrepresented in the list of input SNPs. *snpXplorer* is freely available at https://snpxplorer.net. Source code, documentation, example files and tutorial videos are available within the Help section of *snpXplorer* and at https://github.com/TesiNicco/snpXplorer.

Key points:

- *snpXplorer* shows GWAS summary statistics, regional information and helps deciphering GWAS outcomes
- *snpXplorer* interactively compares association levels of a genomic region across phenotypes
- *snpXplorer* performs variant-to-gene mapping and gene-set enrichment analysis

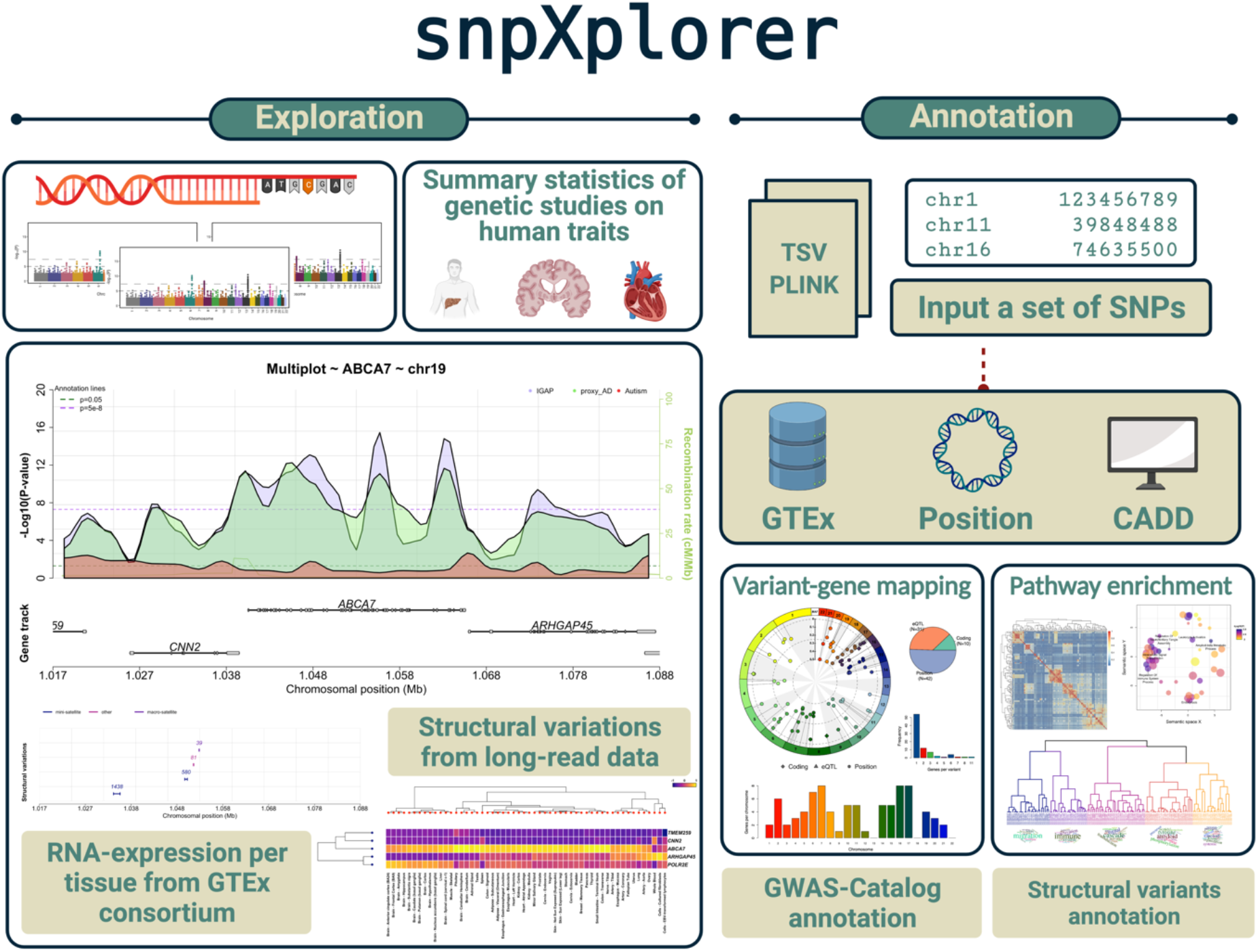

## INTRODUCTION

Genome-wide association studies (GWAS) and sequencing-based association studies are a powerful approach to investigate the genetic basis of complex human phenotypes and their heritability. Facilitated by the cost-effectiveness of both genotyping and sequencing methods and by established analysis guidelines, the number of genetic association studies has risen steeply in the last decade: as of February 2021, the GWAS-Catalog, a database of genetic association studies, contained 4,865 publications and 247,051 variant-trait associations.(1)

To understand how genetic factors affect different traits, it is valuable to explore various annotations of genomic regions as well as how associations relate between different traits. But this requires combining diverse sources of annotation such as observed structural variations (SV), expression-quantitative-trait-loci (eQTL), or chromatin context. Moreover, a framework to quickly visualize and compare association statistics of specific genomic regions across multiple traits is missing, and may be beneficial to the community of researchers working on human genetics. In addition, the functional interpretation of the effects of genetic variants on a gene-, protein- or pathway-level is difficult as often genetic variants lie in non-coding regions of the genome. As a one- to one mapping between genetic variants and affected genes is not trivial in these circumstances, it might be wise to associate multiple genes with a variant. Hence, a profound knowledge of biological databases, bioinformatics tools, and programming skills is often required to interpret GWAS outcomes. Unfortunately, not everyone is equipped with these skills.

To assist human geneticists, we have developed *snpXplorer*, a web-server application written in R that allows (*i*) the rapid exploration of any region in the genome with customizable genomic features, (*ii*) the superimposition of summary statistics from multiple genetic association studies, and (*iii*) the functional annotation and pathway enrichment analysis of SNP sets in an easy-to-use user interface.

## METHODS

### WEB SERVER STRUCTURE

*snpXplorer* is a web-server application based on the R package *shiny* that offers an exploration section and a functional annotation section. The exploration section represents the main interface (Figure 1) and provides an interactive exploration of a (set of) GWAS data sets. The functional annotation section takes as input any list of SNPs, runs a functional annotation and enrichment analysis in the background, and send the results by email.

**Figure 1:**
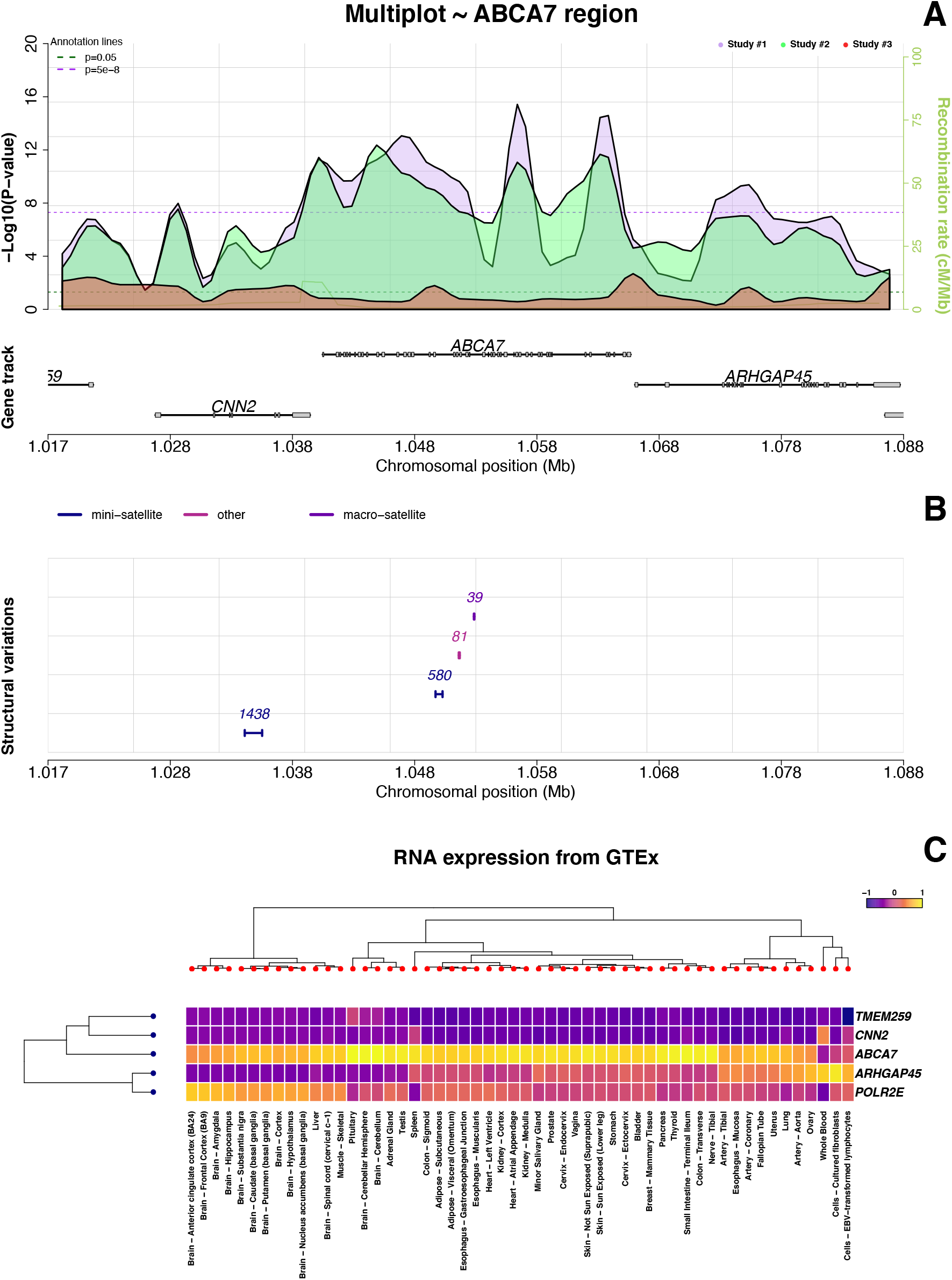
snpXplorer exploration section. **A.** First and main visualisation interface reporting summary statistics of multiple genetic studies as shown with *p*-value profiles. **B.** Structural variants within the region of interest are reported as segments and colored according to their type **C.** Tissue-specific RNA-expression (from Genotype-Tissue-Expression, GTEx) of the genes displayed in the region of interest.

#### Exploration section

First, input data must be chosen, which can either be one of the available summary statistics datasets and/or the user can upload their own association dataset. One of the main novelties in *snpXplorer* is the possibility to select *multiple* association datasets as inputs (including data uploaded by the user). These will be displayed on top of each other with different colours. The available summary statistics will be kept updated. As of February 2021, *snpXplorer* includes genome-wide summary statistics of 23 human traits classified in 5 disease categories: neurological traits (Alzheimer’s disease, family history of Alzheimer’s disease, autism, depression, and ventricular volume),(2–6) cardiovascular traits (coronary artery disease, systolic blood pressure, body-mass index and diabetes),(7–10) immune-related traits (severe COVID infections, Lupus erythematosus, inflammation biomarkers and asthma),(11–14) cancer-related traits (breast, lung, prostate cancers, myeloproliferative neoplasms and Lymphocytic leukaemia),(15–18) and physiological traits (parental longevity, height, education, bone-density and vitamin D intake).(9, 19–22) These summary statistics underwent a process of harmonization: we use the same reference genome (GRCh37, hg19) for all SNP positions, and in case a study was aligned to the GRCh38 (hg38), we translate the coordinates using the liftOver tool.(23) In addition, we only store chromosome, position and *p*-value information for each SNP-association. The user may upload own association statistics to display within *snpXplorer*: the file must have at least chromosome-, position-, and *p*-value columns, and the size should not exceed 600Mb. *snpXplorer* automatically recognizes the different columns, supports PLINK (v1.9+ and v2.0+) association files,(24) and we provide several example files in the Help section of the web-server.

After selecting the input type, the user should set the preferred genome version. By default, GRCh37 is used, however, all available annotation sources are available also for GRCh38, and *snpXplorer* can translate genomic coordinates from one reference version to another. In order to browse the genome, the user can either input a specific genomic position, gene name, variant identifier, or select the scroll option, which allows to interactively browse the genome.

The explorative visualisation consists of 3 separate panels showing (*i*) the SNP summary statistics of the selected input data (Figure 1A), (*ii*) the structural variants in the region of interest (Figure 1B), and (*iii*) the tissue-specific RNA-expression (Figure 1C). The first (and main) visualization panel shows the association statistics of the input data in the region of interest: genomic positions are shown on the x-axis and association significance (in –*log*_10_ scale) is reported on the y-axis. Both the x-axis and the y-axis can be interactively adjusted to extend or contract the genomic window to be displayed. Linkage disequilibrium (LD) patterns are optionally shown for the most significant variant in the region, the input variant, or a different variant of choice. The linkages are calculated using the genotypes of the individuals from the 1000Genome project, with the possibility to select the populations to include.(25) There are two ways to visualise the data: by default, each variant-association is represented as a dot, with dot-sizes optionally reflecting *p*-values. Alternatively, associations can be shown as *p*-value profiles: to do so, (*i*) the selected region is divided in bins, (*ii*) a local maximum is found in each bin based on association *p-*value, and (*iii*) a polynomial regression model is fitted to the data, using the *p-*value of all local maximum points as dependent variable and their genomic position as predictors. Regression parameters, including the number of bins and the smoothing value, can be adjusted. Gene names from RefSeq (v98) are always adapted to the plotted region.(26) Finally, recombination rates from HapMap II, which give information about recombination frequency during meiosis, are optionally shown in the main plot interface.(27)

The second panel shows structural variations (SV) in the region of interest. These are extracted from three studies that represent the state-of-the-art regarding the estimation of major structural variations across the genome using third-generation sequencing technologies (*i.e.* long read sequencing).(28–30) Structural variations are represented as segments: the size of the segment codes for the maximum difference in allele sizes of the SVs as observed in the selected studies. Depending on the different studies, structural variations are annotated as insertions, deletions, inversions, copy number alterations, duplications, mini-, micro- and macro-satellites, and mobile element insertions (Alu elements, LINE1 elements, and SVAs).

The third panel shows tissue-specific RNA-expression (from the Genotype-Tissue-expression consortium, GTEx) of the genes displayed in the selected genomic window.(31) The expression of these genes across 54 human tissues is scaled and reported as a heatmap. Hierarchical clustering is applied on both the genes and the tissues, and the relative dendrograms are reported on the sides of the heatmap.

The side panel allows the user to interact with the exploration section. In order to guide the user through all the available inputs and options, help messages automatically appear upon hovering over items. The side panel reports (*i*) the top 10 variants with highest significance (together with the trait they belong to, in case multiple studies were selected), and (*ii*) the top eQTLs associations (by default, eQTLs in blood are shown, and this can be optionally changed), and cross-references including GeneCards, GWAS-catalog, and LD-hub.(1, 32, 33) Finally, download buttons allow to download a high-quality image of the different visualisation panels as well as the tables reporting the top SNP and eQTL associations, the SVs in the selected genomic window, and the LD table.

#### Functional annotation section

The functional annotation pipeline consists of a two-step procedure: firstly, genetic variants are linked to likely affected genes (*variant-gene mapping*); and, secondly, the likely affected genes are tested for pathway enrichment (*gene-pathway mapping*). In the *variant-gene mapping*, genetic variants are linked to the most likely affected gene(s) by (*i*) associating a variant to a gene when the variant is annotated to be coding by the Combined Annotation Dependent Depletion (CADD, v1.3), (*ii*) annotating a variant to genes based on found expression-quantitative-trait-loci (eQTL) from GTEx (v8, with possibility to choose the tissue(s) of interest), or (*iii*) mapping a variant to genes that are within distance *d* from the variant position, starting with *d* ≤ 50*kb*, up to *d* ≤ 500*kb*, increasing by 50*kb* until at least one match is found (from RefSeq v98).(26, 31, 34) Note that this procedure might map multiple genes to a single variant, depending on the effect and position of each variant.

Then, we first report whether the input SNPs as well as their likely associated genes were previously associated with any trait in the GWAS-Catalog (traits are coded by their Experimental Factor Ontology (EFO) term). For this analysis, we downloaded all significant SNP-trait associations of all studies available in the GWAS-Catalog (v1.0.2, available at https://www.ebi.ac.uk/gwas/docs/file-downloads), which includes associations with *p*<9×10^−6^. Given a set of input SNPs associated with a set of genes, this analysis results in a set of traits (provided that the SNPs and/or the genes were previously associated with a trait). Hereto, we plot the number of SNPs in the list of uploaded SNPs that associate with the trait (expressed as a fraction). To correct for multiple genes being associated with a single variant, we estimate these fractions by sampling (500 iterations) one gene from the pool of genes associated with each variant, and averaging the resulting fractions across the sampling. Summary tables of the GWAS-Catalog analysis, including also EFO URI links for cross-referencing are provided as additional output.

Next, we report on the structural variations that lie in the vicinity (10*kb* upstream and downstream) of the input SNPs, and present information such as SV start and end position, SV type, maximum difference in allele size, and genes likely associated with the relative SNPs.

Finally, we perform a gene-set enrichment analysis to find molecular pathways enriched within the set of genes associated with the input variants. Also, here we use the mentioned sampling technique to avoid a potential enrichment bias due to multiple genes being mapped to the same variant (this time the sampling is used to calculate *p-*values for each term). The gene-set enrichment analysis is performed using the *Gost* function from the R package *gprofiler2*.(35) The user can specify several gene-set sources, such as Gene Ontology (release 2020-12-08),(36) KEGG (release 2020-12-14),(37) Reactome (release 2020-12-15),(38) and Wiki-pathways (release 2020-12-10).(39) The full table of the gene-set enrichment analysis comprising all tested terms and their relative sampling-based *p*-values is sent to the user.

For each of the selected gene-set sources, the significant enriched terms are plotted (up to FDR<10%). In case the Gene Ontology is chosen as gene-set source, we additionally reduce the visual complexity of the enriched biological processes using (*i*) the REVIGO tool and (*ii*) a term-based clustering approach.(40) We do so because the interpretation of gene-set enrichment analyses is typically difficult due to the large number of terms. Clustering enriched terms then helps to get an overview, and thus eases the interpretation of the results. Briefly, REVIGO masks redundant terms based on a semantic similarity measure, and displays enrichment results in an embedded space via eigenvalue decomposition of the pairwise distance matrix. In addition to REVIGO, we developed a term-based clustering approach to remove redundancy between enriched terms. To do so, we first calculate a semantic similarity matrix between all enriched terms, and then apply hierarchical clustering on the obtained distance matrix. We estimate the optimal number of clusters using a dynamic cut tree algorithm and plot the most recurring words of the terms underlying each cluster using wordclouds. We use *Lin* as semantic distance measure for both REVIGO and our term-based clustering approach.(41, 42) Figures representing REVIGO results, the semantic similarity heatmap (showing relationships between enriched terms), the hierarchical clustering dendrogram, and the wordclouds of each clusters, are generated. Finally, all tables describing REVIGO analysis and our term-based clustering approach (including all enriched terms and their clustering scheme) are produced and sent as additional output to the user for further manipulation. Note that the initial significant GO terms are not removed and also included in the reporting.

## RESULTS

### Case Study

To illustrate the performances of *snpXplorer*, we explored the most recent set of common SNPs associated with late-onset Alzheimer’s disease (AD, N=83 SNPs, Table S1).(43) Using this dataset as case study, we show the benefits of using *snpXplorer* in a typical scenario. Briefly, AD is the most prevalent type of dementia at old age, and is associated with a progressive loss of cognitive functions, ultimately leading to death. In its most common form (late-onset AD, with age at onset typically >65 years), the disease is estimated to be 60-80% heritable. With an attributable risk of ~30%, genetic variants in *APOE* gene represent the largest common genetic risk factor for AD. In addition to *APOE,* the genetic landscape of AD now counts 83 common variants that are associated with a slight modification of the risk of AD. Understanding the genes most likely involved in AD pathogenesis as well as the crucial biological pathways is warranted for the development of novel therapeutic strategies for AD patients.

We retrieved the list of AD-associated genetic variants in Table 1 of the preprint from *Bellenguez et al*, 2020.(43) This study represent the largest GWAS on AD performed to date, and resulted in 42 novel SNPs reaching genome-wide evidence of association with AD. The exploration section of *snpXplorer* can be firstly used to inspect the association statistics of the novel SNP-associations in previous studies of the same trait (*i.e.* International Genomics of Alzheimer Project (IGAP) and family history of AD (proxy_AD)). Specifically, a suggestive degree of association in these regions is expected to be found in earlier studies. As expected, suggestive association signals were already observed for the novel SNPs, increasing the likelihood that these novel SNPs are true associations (Figure S1).

After the first explorative analysis, we pasted the variant identifiers (rsIDs) in the annotation section of *snpXplorer*, specifying rsid as input type, Gene Ontology and Reactome as gene-sets for the enrichment analysis, and Blood as GTEx tissue for eQTL (*i.e.* the default value). The N=83 variants were linked to a total of 162 genes, with N=54 variants mapping to 1 gene, N=12 variants mapping to 2 genes, N=7 variants mapping to 3 genes, N=2 variants mapping to 4 genes, N=1 variant mapping to 5 genes, N=4 variants mapping to 4 genes, and N=1 variant mapping to 7, 8 and 11 genes (Figure S2). N=10 variants were found to be coding variants, N=31 variants were found to be eQTL, and N=42 variants were annotated based on their genomic position. These results are returned to the user in the form of a (human and machine-readable) table, but also in the form of a summary plot (Figure 2A and Figure S2). These graphs not only inform the user about the effect of the SNPs of interest (for example, a direct consequence on the protein sequence in case of coding SNPs, or a regulatory effect in case of eQTLs or intergenic SNPs), but also suggest the presence of more complex regions: for example, Figure S2B indicates the number of genes associated with each SNP, which normally increases for complex, gene-dense regions such as HLA-region or IGH-region.

**Figure 2:**
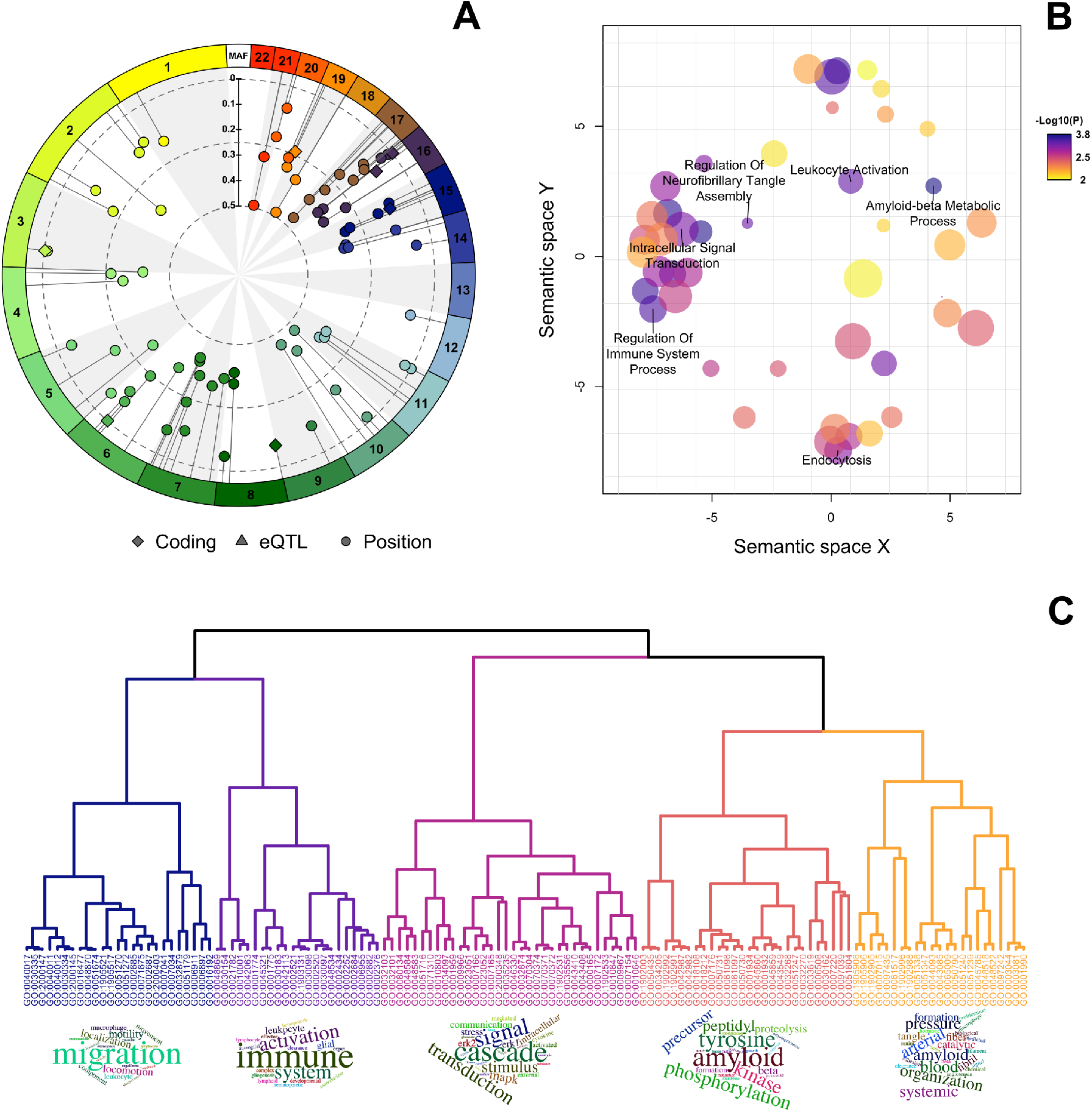
Results of the functional annotation of N=83 variants associated with Alzheimer’s disease (AD). **A.** The circular summary figure shows the type of annotation of each genetic variant used as input (coding, eQTL or annotated by their positions) as well as each variant’s minor allele frequency and chromosomal distribution. **B.** REVIGO plot, showing the remaining GO terms after removing redundancy based on a semantic similarity measure. The colour of each dot codes for the significance (the darker, the more significant), while the size of the dot codes for the number of similar terms removed from REVIGO. **C.** Results of our term-based clustering approach. We used Lin as semantic similarity measure to calculate similarity between all GO terms. We then used ward-d2 as clustering algorithm, and a dynamic cut tree algorithm to highlight clusters. Finally, for each cluster we generated wordclouds of the most frequent words describing each cluster.

In order to prioritize candidate genes, the authors of the original publication integrated (*i*) eQTLs and colocalization (eQTL coloc) analyses combined with expression transcriptome-wide association studies (eTWAS) in AD-relevant brain regions; (*ii*) splicing quantitative trait loci (sQTLs) and colocalization (sQTL coloc) analyses combined with splicing transcriptome-wide association studies (sTWAS) in AD-relevant brain regions; (*iii*) genetic-driven methylation as a biological mediator of genetic signals in blood (MetaMeth).(43) In order to compare the SNP-gene annotation of the original study with that of *snpXplorer,* we counted the total number of unique genes associated with the SNPs (*i*) in the original study (N=97), (*ii*) using our annotation procedure (N=136), and (*iii*) the intersection between these gene sets (N=79). When doing so, we excluded regions mapping to the *HLA*-gene cluster and *IGH*-gene clusters (3 SNPs in total) as the original study did not report gene names but rather *HLA*-cluster and *IGH*-cluster. Nevertheless, our annotation procedure correctly assigned *HLA*-related genes and *IGH*-related genes with these SNPs. The number of intersecting genes was significantly higher than what could be expected by chance (*p*=0.03, based on one-tail *p-*value of binomial test, Table S2). For 6 SNPs, the gene annotated by our procedure did not match the gene assigned in the original study. Specifically, for 4/6 of these SNPs, we found significant eQTLs in blood (rs60755019 with *ADCY10P1*, rs7384878 with *PILRB*, *STAG3L5P*, *PMS2P1*, *GIGYF1*, and *EPHB4* genes, rs56407236 with *FAM157C* gene, and rs2526377 with *TRIM37* gene), while the original study reported the closest genes as most likely gene (rs60755019 with *TREML2* gene, rs7384878 with *SPDYE3* gene, rs56407236 with *PRDM7* gene and rs2526377 with *TSOAP1* gene). In addition, we annotated SNPs rs76928645 and rs139643391 to *SEC61G* and *WDR12* genes (closest genes), while the original study, using eQTL and TWAS in AD-relevant brain regions, annotated these SNPs to *EGFR* and *ICA1L*/*CARF* genes. While the latter two SNPs were likely mis-annotated in our procedure (due to specific datasets used for the annotation), our annotation of the former 4 SNPs seemed robust, and further studies will have to clarify the annotation of these SNPs.

With the resulting list of input SNPs and (likely) associated genes, we probed the GWAS-Catalog and the datasets of structural variations for previously reported associations. We found a marked enrichment in the GWAS-Catalog for Alzheimer’s disease, family history of Alzheimer’s disease, and lipoprotein measurement (Figure S3, Table S3 and Table S4). The results of this analysis are relevant to the user as they indicate other traits that were previously associated with the input SNPs. As such, they may suggest relationships between different traits, for example in our case study they suggest the involvement of cholesterol and lipid metabolism in AD, a known relationship.(44) Next, we searched for all structural variations in a region of 10kb surrounding the input SNPs, and we found that for 39/83 SNPs, a larger structural variations was present in the vicinity (Table S5), including the known VNTR (variable number of tandem repeats) in *ABCA7* gene,(45) and the known CNV (copy number variation) in *CR1, HLA-DRA,* and *PICALM* genes (Table S5).(46–48) This information may be particularly interesting for experimental researchers investigating the functional effect of SVs, and could be used to prioritize certain genomic regions. Because of the complex nature of large SVs, these regions have been largely unexplored, however technological improvements now make it possible to accurately measure SV alleles.

We then performed our (sampling-based) gene-set enrichment analysis using Gene Ontology Biological Processes (GO:BP, default setting) and Reactome as gene-set sources, and Blood as tissue for the eQTL analysis. After averaging *p*-values across the number of iterations, we found N=132 significant pathways from Gene Ontology (FDR<1%) and N=4 significant pathways from Reactome (FDR<10%) (Figure S4 and Table S6). To facilitate the interpretation of the gene-set enrichment results, we clustered the significantly enriched terms from Gene Ontology based on a semantic similarity measure using REVIGO (Figure 2B) and our term-based clustering approach (Figure 2C). Both methods are useful as they provide an overview of the most relevant biological processes associated with the input SNPs. Our clustering approach found five main clusters of GO terms (Figure 2C and Figure S5). We generated wordclouds to guide the interpretation of the set of GO terms of each cluster (Figure 2C). The five clusters were characterized by (1) trafficking and migration at the level of immune cells, (2) activation of immune response, (3) organization and metabolic processes, (4) beta-amyloid metabolism and (5) amyloid and neurofibrillary tangles formation and clearance (Figure 2C). All these processes are known to occur in the pathogenesis of Alzheimer’s disease from other previous studies.(43, 44, 49, 50) We observed that clusters generated by REVIGO are more conservative (*i.e.* only terms with a high similarity degree were merged) as compared to our term-based clustering which generates a higher-level overview. In the original study (Table S15 from (43)), the most significant gene sets related to amyloid and tau metabolism, lipid metabolism and immunity. In order to calculate the extent of term overlap between results from the original study and our approach, we calculated semantic similarity between all pairs of significantly enriched terms in both studies. In addition to showing pairwise similarities between all terms, this analysis also shows how the enriched terms in the original study relate to the clusters found using our term-based approach. We observed patterns of high similarity between the significant terms in both studies (Figure S6). For example, terms in the “Activation of immune system” and the “Beta-amyloid metabolism” clusters (defined with our term-based approach), reported high similarities with specific subsets of terms from the original study. This was expected as these clusters represent the most established biological pathways associated with AD. The cluster “Trafficking of immune cells” had high similarity with a specific subset of terms from the original study, yet we also observed similarities with the “Activation of immune system” cluster, in agreement with the fact that these clusters were relatively close also in tree structure (Figure 2C). Similarly, high similarities were observed between the “Beta-amyloid metabolism” and the “Amyloid formation and clearance” clusters. Finally, the “Metabolic processes” had high degree of similarity with a specific subset of terms, but also with terms related to “Activation of immune system” cluster. Altogether, we showed that (*i*) enriched terms from the original study and our study had a high degree of similarity, and (*ii*) that the enriched terms of the original study resembled the structure of our clustering approach. The complete analysis of 83 genetic variants took about 30 minutes to complete.

## DISCUSSION

Despite the fact that many summary statistics of genetic studies have been publicly released, the integration of such a large amount of data is often difficult and requires specific tools and knowledge. Even simple tasks, such as the rapid interrogation of how well a certain genomic region associates with a specific trait or multiple traits can be frustrating and time consuming. Our main objective to develop *snpXplorer* was the need for an easy-to-use and user-friendly framework to explore, analyse and integrate outcomes of GWAS and other genetic studies. *snpXplorer* showed to be a robust tool that can support a complete GWAS analysis, from the exploration of specific regions of interest to the variant-to-gene annotation, gene-set enrichment analysis and interpretation of associated biological pathways.

To our knowledge, the only existing web-server that offers a similar explorative framework as *snpXplorer* is the GWAS-Atlas.(51) GWAS-Atlas was primarily developed as a database of publicly available GWAS summary statistics. It offers possibilities to visualise Manhattan and quantile-quantile (QQ) plots, to perform downstream analyses using MAGMA statistical framework, and to study genetic correlation between traits by means of LD score regression.(52, 53) However, *snpXplorer* was developed mainly for visualisation purposes, and thus incorporate multiple unique features such as the possibility to visualise multiple GWAS datasets simultaneously or to upload an external association dataset for additional comparisons with existing datasets. Moreover, *snpXplorer* annotates these visualisations with several genomic features such as structural variations, recombination rates, LD patterns and eQTLs. All the relevant information showed in *snpXplorer*, such as top SNP information, eQTL tables, LD tables and structural variants can be easily downloaded for further investigations. Further, we would like to stress the relevance of overlaying the GWAS results with structural variants found by third-generation sequencing. Such structural variations have already been shown to play a significant role for several traits, in particular for neurodegenerative diseases, and *snpXplorer* is thus far the only web-server where such information can be visualized in the context of GWAS summary statistics.(45, 46, 54, 55)

We do acknowledge that for an in-depth functional annotation analysis of GWAS, the possibility of integrating additional ad-hoc information (such as eQTLs, sQTLs, eTWAS and sTWAS from specific disease-related regions) may improve the analysis, but such data is not always available, is time consuming and requires deep knowledge. Several online and offline tools have been developed with a similar goal, *e.g.* SNPnexus, ANNOVAR, FUMA and Ensembl VEP.(56–59) Some of these tools are characterized by a larger list of annotation sources, for example implementing multiple tools for variant effect prediction (*e.g.* SNPnexus, Ensembl VEP or ANNOVAR), or more extensive pathway enrichment analyses at the tissue- and cell-type level (*e.g.* FUMA). We have shown that *snpXplorer* provides similar results in terms of annotation capabilities and gene-set enrichment analysis as compared to existing tools. Yet, *snpXplorer* has several unique features for the functional annotation section, such as the extensive interpretation analysis implemented in REVIGO, our term-based clustering approach and the wordcloud visualisation, or the possibility to associate multiple genes with each SNP during gene-set enrichment analysis. Moreover, *snpXplorer* development will continue by implementation of additional annotation sources and analyses. Altogether, we showed that *snpXplorer* is a promising functional annotation tool to support a typical GWAS analysis. As such, it has been previously applied for the annotation and downstream analysis of genetic variants associated with Alzheimer’s disease and human longevity.(42, 60)

### Future updates

For future updates, we plan to keep updated and increase the list of summary statistics available to be displayed in the exploration section. In its current version, the exploration section of *snpXplorer* requires the user to define a region of interest to look, while genome-wide comparisons are not considered. However, it is our intention to implement a genome-wide comparison across GWAS studies that, given a set of input GWASs and a significance threshold *alpha*, reports all SNPs with a *p*<*alpha* across the studies, allowing for a more rapid visualisation of overlapping SNP-associations. Moreover, we plan to increase the number of annotation sources and available options in the annotation section (for example, including methylation-QTL, protein-QTL and splicing-QTL). Finally, we are also working towards adding a framework to calculate weighted polygenic risk scores given a set of individuals’ genotypes and a reference study to take variant effect-sizes from.

## AVAILABILITY

*snpXplorer* is an open-source web-server available at https://snpxplorer.net. Tutorial videos, full documentation and link to code are available in the *Help* page of the web-server. *snpXplorer* is running as from March-2020, was tested both within and outside our group, and runs steadily on both Unix and Windows most common browsers (Safari, Google Chrome, Microsoft Edge, Internet Explorer, and Firefox). For certain steps, *snpXplorer* does rely on external tools and sources (e.g. REVIGO), and consequently depends on their availability. Although discouraged, the tool can also be installed locally on your machine: additional information on how to do it are available in our *github* at https://github.com/TesiNicco/snpXplorer, however, we note that for the stand-alone version additional files should be downloaded separately, for example, all summary statistics. *snpXplorer* requires R (v3.5+) and python (v3+) correctly installed and accessible in your system. *snpXplorer* uses the following R packages: *shiny, data.table, stringr, ggplot2, liftOver, colourpicker, rvest, plotrix, parallel, SNPlocs.Hsapiens.dfSNP144.GRCh37, lme4, ggsci, RColorBrewer, gprofiler2, GOSemSim, GO.db, org.Hs.eg.db, pheatmap, circlize, devtools, treemap, basicPlotteR, gwascat, GenomicRanges, rtracklayer, Homo.sapiens, BiocGenerics*, and the following python libraries: *re, werkzeug, robobrowser, pygosemsim, numpy, csv, networkx* and *sys*.

## ACKNOWLEDGEMENTS AND FUNDING

The authors declare no conflict of interests. This work was supported by Stichting Alzheimer Nederland (WE09.2014-03), Stichting Diorapthe, horstingstuit foundation, Memorabel (ZonMW projectnumber 733050814) and Stichting VUmc Fonds.

