## Supplementary Figures for "snpXplorer: a web application to explore human SNP-associations and annotate SNP-sets"

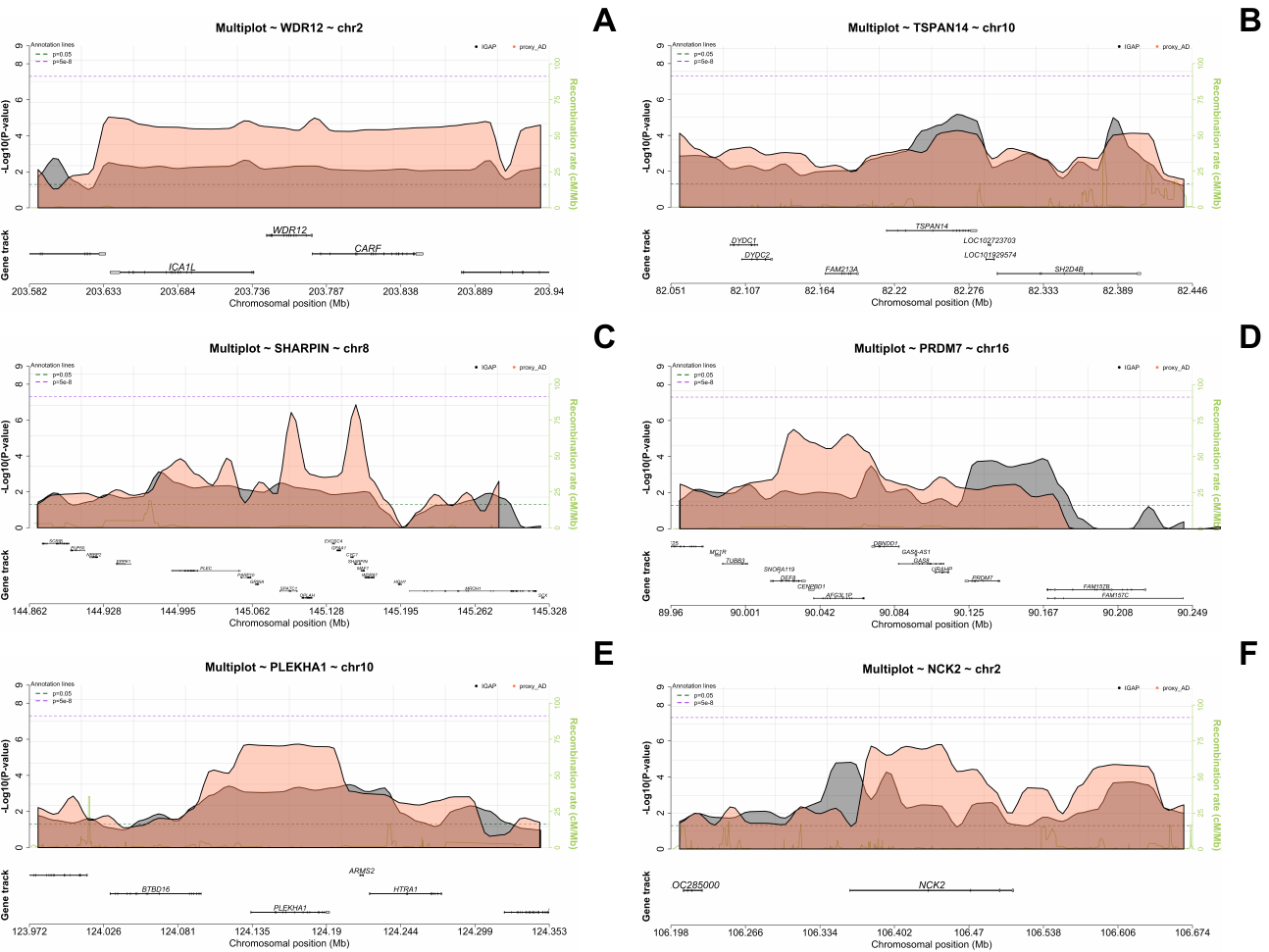

**Figure S1: Rapid exploration of novel SNP-associations in existing GWAS datasets. A.** The figure shows the region surrounding *WDR12* gene, for which a novel SNP-association was found in the case-study GWAS of Alzheimer’s disease (AD). Two previous studies of AD are plotted, and show that suggestive association signals were present in earlier studies, yet the association did not reach genome-wide statistical significance, likely due to sample size. Similar plots show the regions surrounding *TSPAN14* (**B**), *SHARPIN* (**C**), *PRDM7* (**D**) , *PLEKHA1* (**E**) and *NCK2* (**F**).

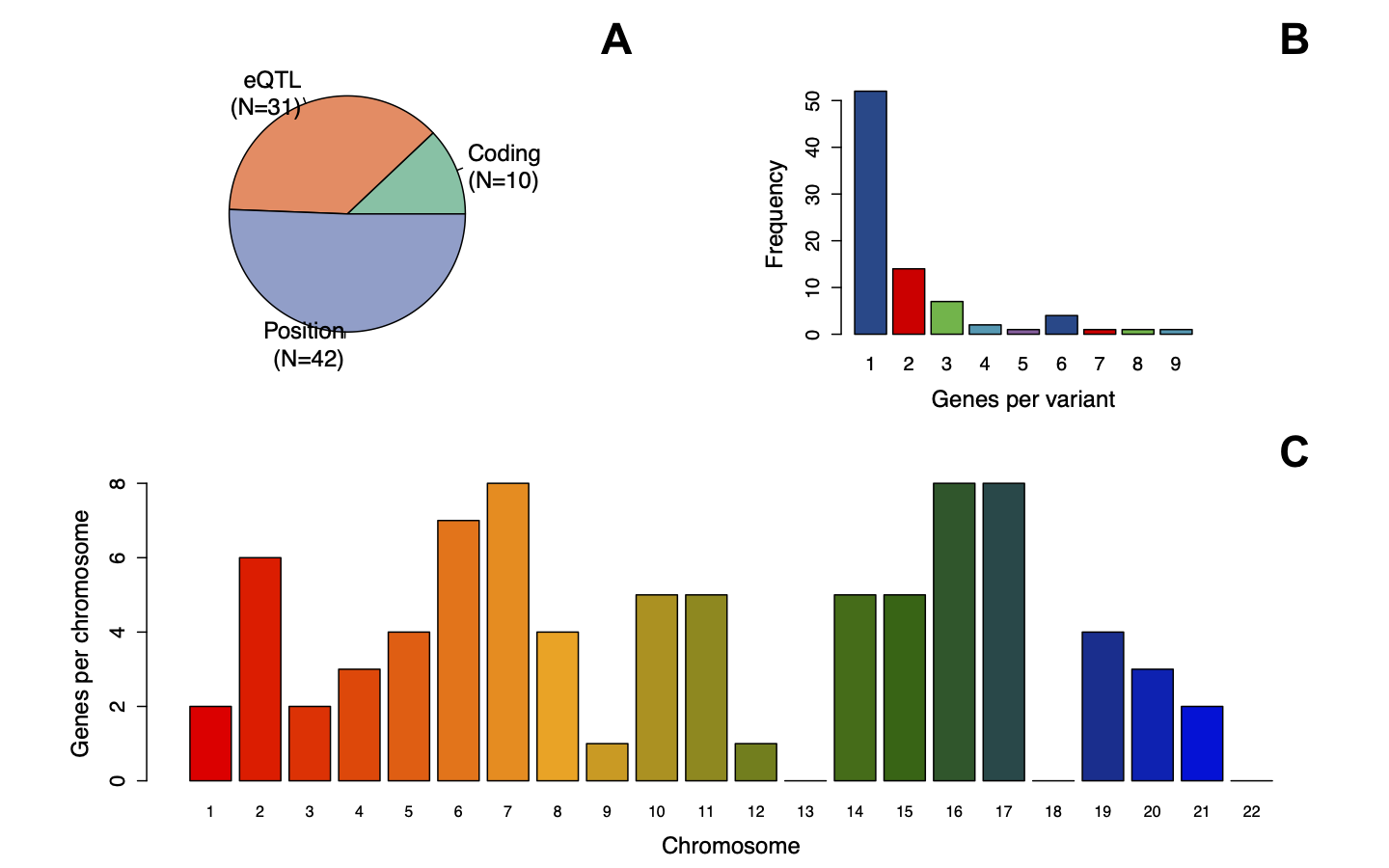

**Figure S2: Variant-gene mapping procedure. The summary figure shows the type of genetic variants used as input (pie plot), classified as coding, eQTL or annotated by their positions.** Top-right barplot shows the number of genes associated with each variant. The central barplot shows the chromosomal distribution of all input variants. The circular summary shows the frequency, the type and the chromosomal distribution of all input variants. Supplementary Figure 1: Variant-gene mapping procedure. The summary figure shows the type of genetic variants used as input (top-left plot), classified as coding, eQTL or annotated by their positions. Top-right barplot shows the number of genes associated with each variant. The central barplot shows the chromosomal distribution of all input variants. The circular summary shows the frequency, the type and the chromosomal distribution of all input variants.

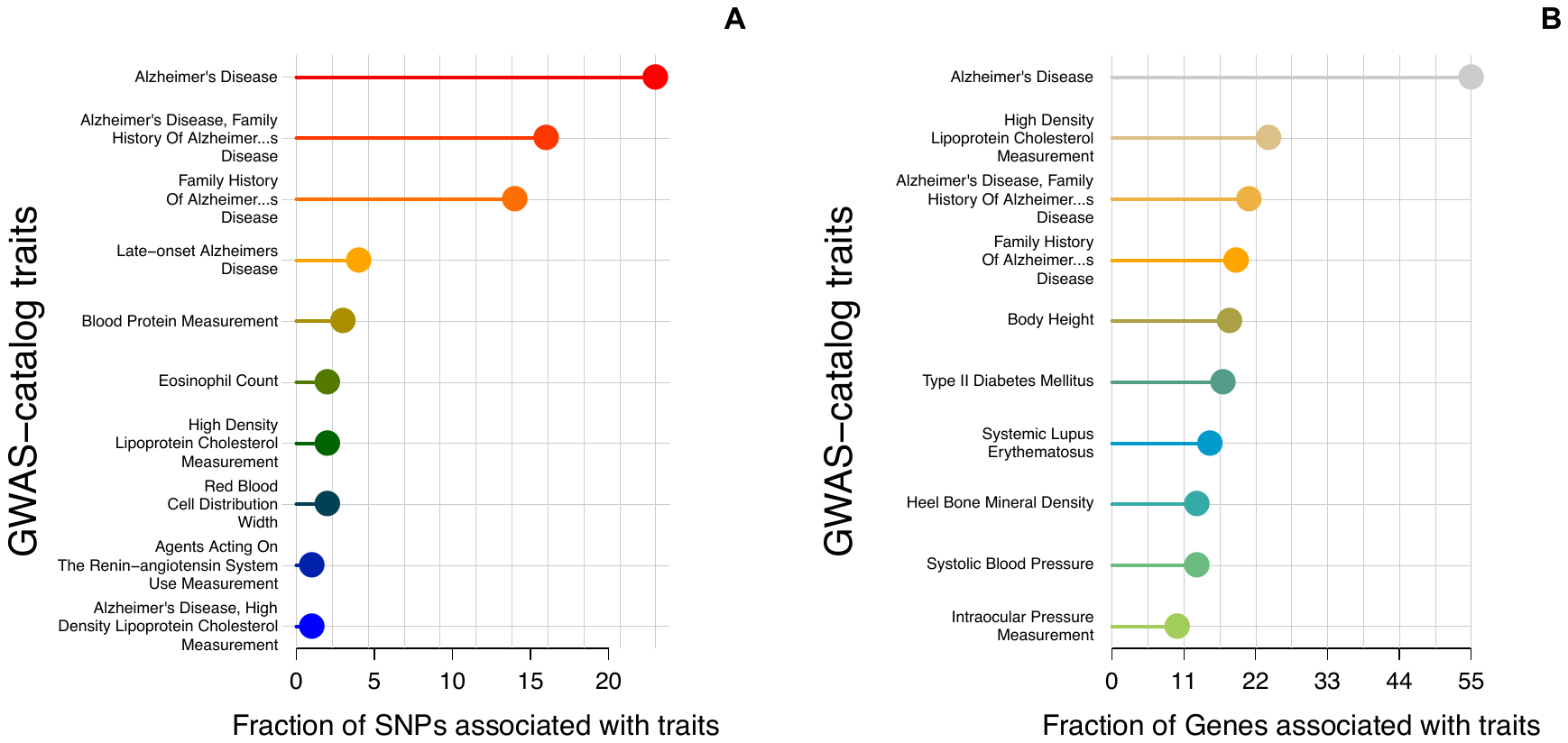

**Figure S3: Fraction of SNPs and Genes association with traits in the GWAS Catalog. A.** Number of input SNPs previously associated with traits in the GWAS catalog. **B.** Fraction of genes (associated with input SNPs) previously associated with traits in the GWAS Catalog.

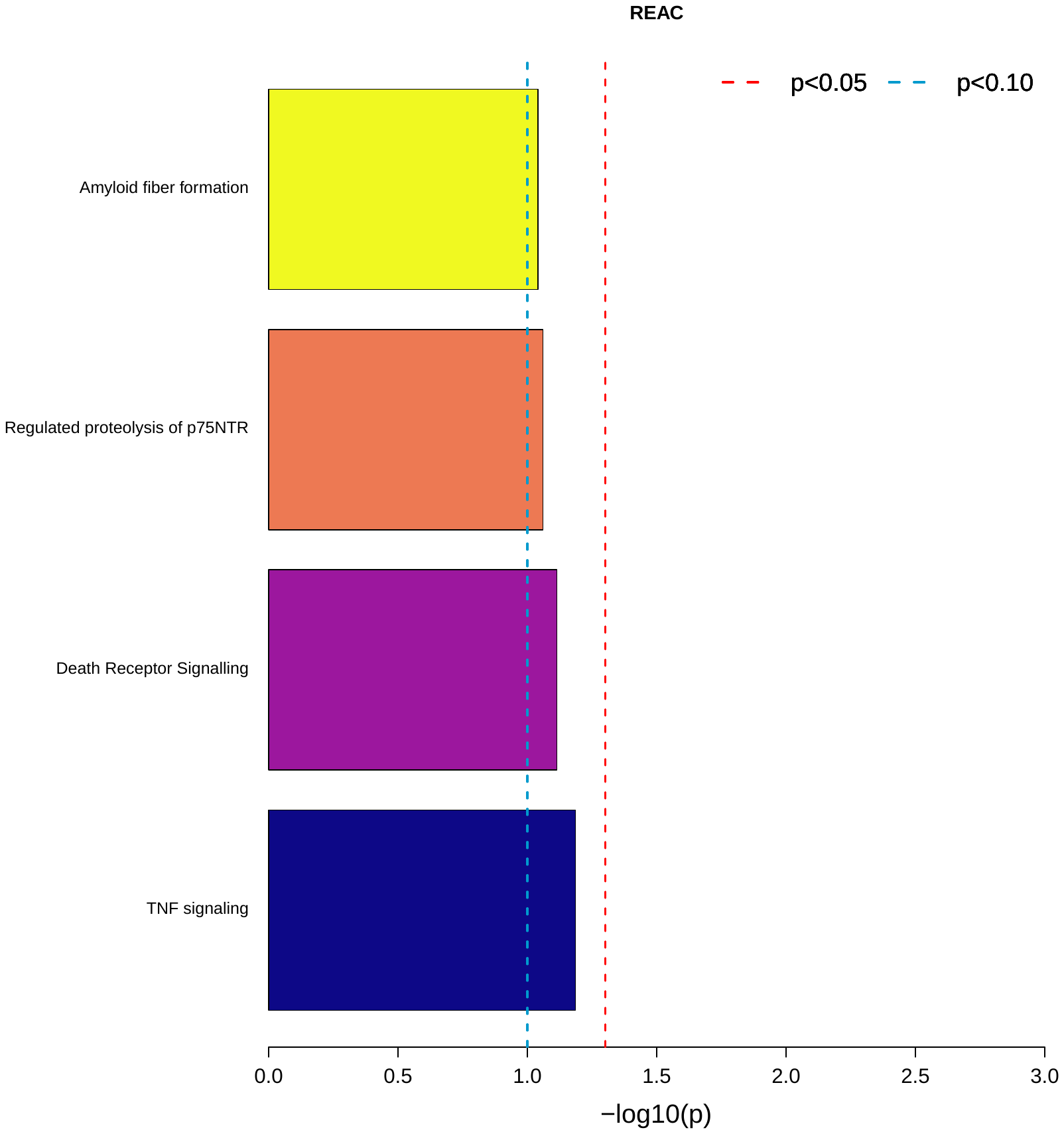

**Figure S4: Gene-set enrichment analysis.** The figure shows the barplot of the most significant pathways (FDR<10%) from Reactome.

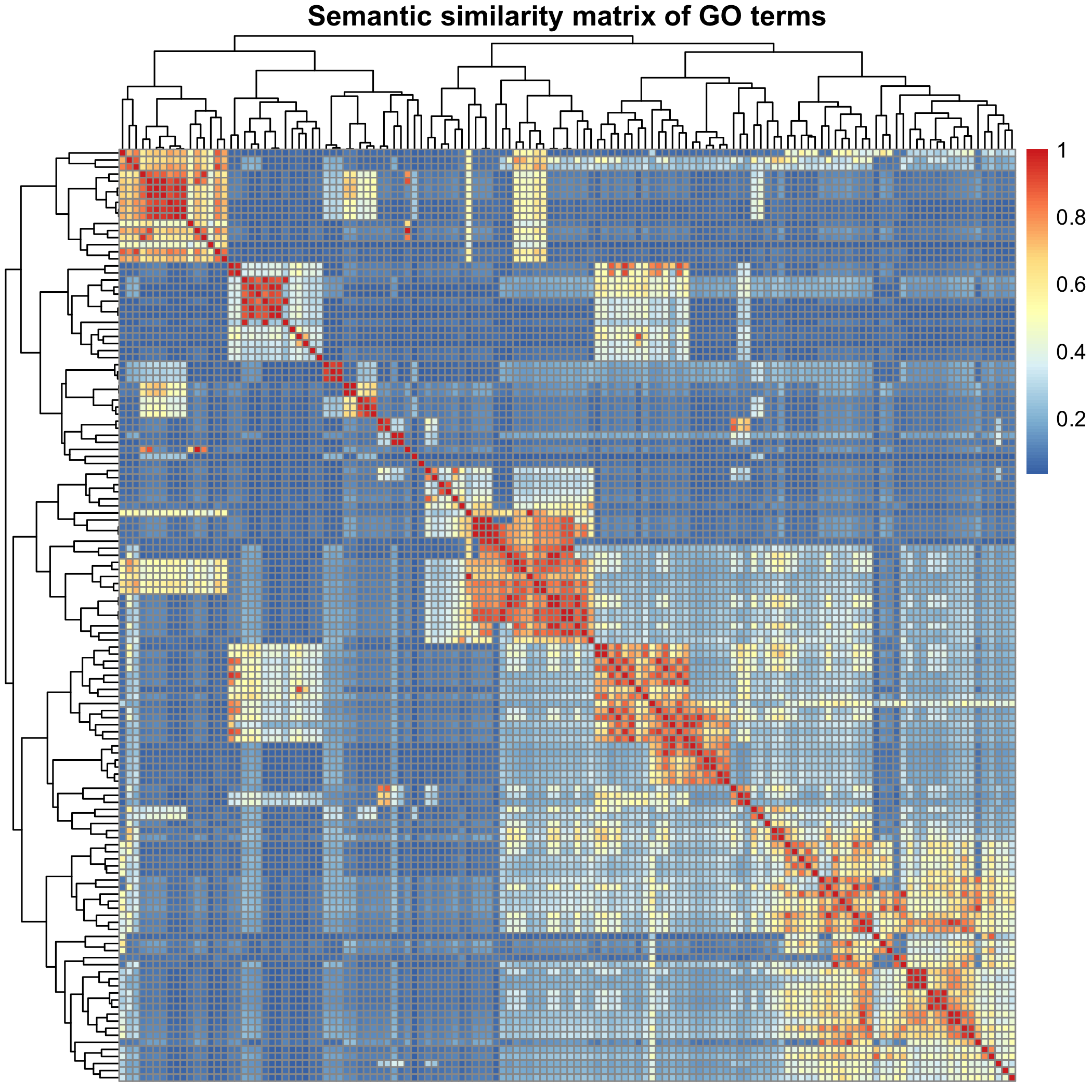

**Figure S5: Semantic similarity matrix.** The plot shows the semantic similarity matrix between all significantly enriched GO (Gene Ontology) biological processes terms (N=132). As semantic similarity, we used Lin.

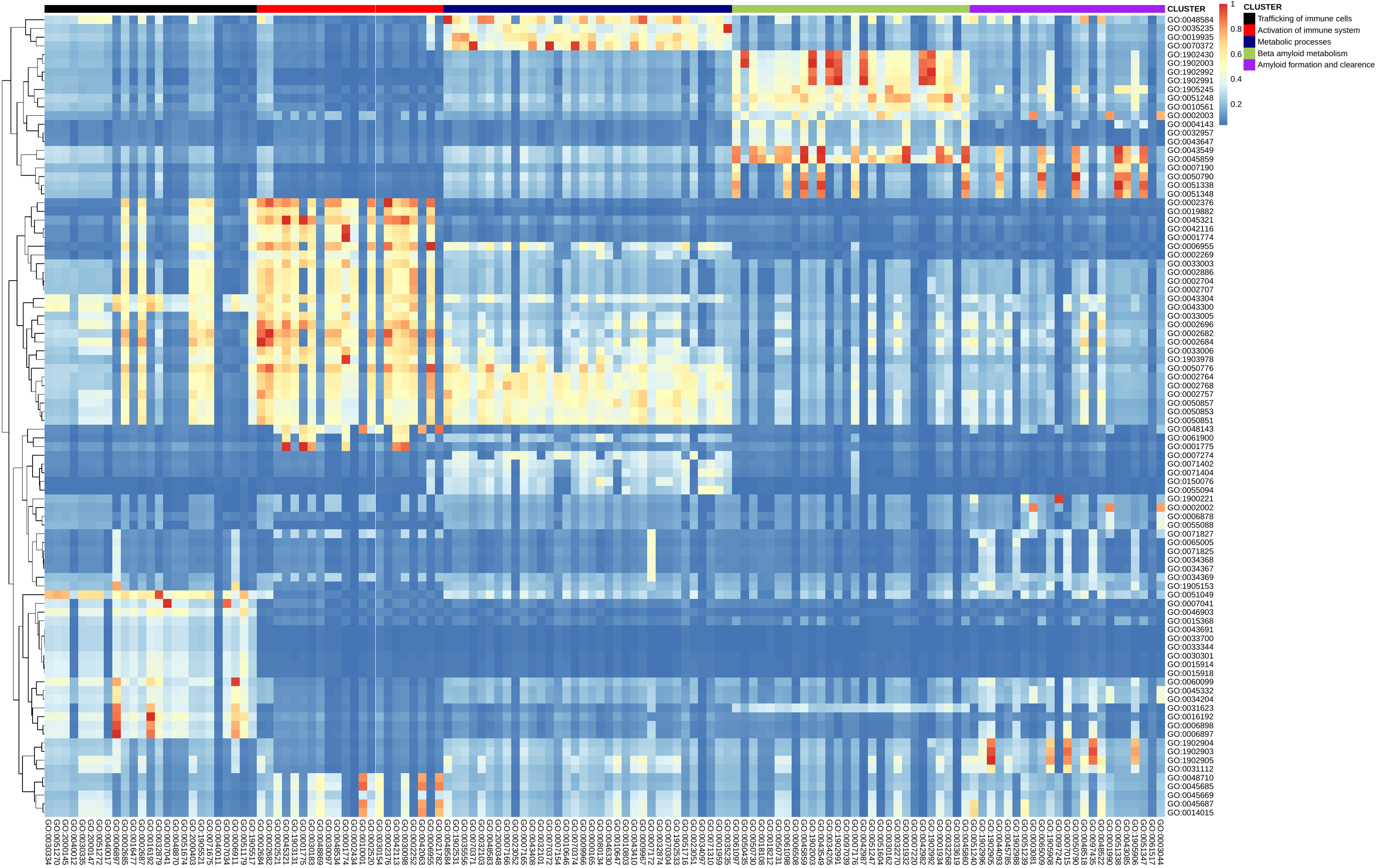

**Figure S6: Comparison of gene-set enrichment results in the original study and using *snpXplorer* functional annotation section.** The heatmap shows the pairwise semantic similarity values between all significantly enriched terms in the original study (y-axis, N=92) and using *snpXplorer* functional annotation section (x-axis, N=132). The terms from our study (x-axis) are ordered based on their assigned cluster as a result of our term-based clustering approach. Large similarity patterns are visible in the heatmap, especially of terms (from the original study) mapping to “Activation of immune response” cluster (red cluster) and to “Beta-amyloid metabolism” cluster (green cluster). Some enriched terms mapping to “Trafficking of immune cells” (black cluster) had high similarity with “Activation of immune response” (purple cluster) cluster, and some terms mapping to “Amyloid formation and clearance” had high similarity with “Beta-amyloid metabolism”, resembling the structure of the tree constructed in our study (Figure 2C). The remaining “Metabolic processes” cluster (blue cluster) had high similarity with a specific subset of enriched terms, but we also observed high similarity with the “Activation of immune system” cluster.
